# Neural Circuit Dynamics for Sensory Detection

**DOI:** 10.1101/2020.01.28.923839

**Authors:** Sruti Mallik, Srinath Nizampatnam, Anirban Nandi, Debajit Saha, Baranidharan Raman, ShiNung Ching

**Affiliations:** Dept. of Electrical and Systems Engineering, Washington University in St. Louis, St. Louis, MO 63130, USA; Allen Institute of Brain Science, Seattle, WA 98109, USA; Dept. of Biomedical Engineering, Michigan State University, East Lansing, MI 48824, USA; Dept. of Biomedical Engineering, Washington University in St. Louis, MO 63130, USA

## Abstract

We consider the question of how sensory networks enable the detection of sensory stimuli in a combinatorial coding space. We are specifically interested in the olfactory system, wherein recent experimental studies have reported the existence of rich, enigmatic response patterns associated with stimulus onset and offset. This study aims to identify the functional relevance of such response patterns, i.e., what benefits does such neural activity provide in the context of detecting stimuli in a natural environment. We study this problem through the lens of normative, optimization-based modeling. Here, we define the notion of a low dimensional latent representation of stimulus identity, which is generated through action of the sensory network. The objective of our optimization framework is to ensure high fidelity tracking of a nominal representation in this latent space in an energy efficient manner. It turns out that the optimal motifs emerging from this framework possess morphological similarity with prototypical onset and offset responses observed *in vivo*. Furthermore, this objective can be exactly achieved by a network with reciprocal excitatory-inhibitory competitive dynamics, similar to interactions between principal neurons (PNs) and local neurons (LNs) in the early olfactory system of insects. The derived model also makes several predictions regarding maintenance of robust latent representations in the presence of confounding background information and tradeoffs between the energy of sensory activity and resultant behavioral measures such as speed and accuracy of stimulus detection.

**Significance Statement:** A key area of study in olfactory coding involves understanding the transformation from high-dimensional sensory stimulus to low-dimensional decoded representation. Here, we treat not only the dimensionality reduction of this mapping but also its temporal dynamics, with specific focus on stimuli that are temporally continuous. We examine through optimization-based synthesis how sensory networks can track representations without prior assumption of discrete trial structure. We show that such tracking can be achieved by canonical network architectures and dynamics, and that the resulting responses resemble observations from neurons in the insect olfactory system. Thus, our results provide hypotheses regarding the functional role of olfactory circuit activity at both single neuronal and population scales.

## Introduction

We consider the question of how early sensory networks produce actionable neural representation in response to sensory stimuli arriving in a dynamic fashion. The specific focus here is on the olfactory system wherein the architecture is schematically conserved across species (Strausfeld and Hildebrand, 1999). In this system, early networks receive external excitation from periphery and transform it into intermediate representations which are then routed to higher-level brain areas for further processing and behavioral and motor response generation (Kay and Stopfer, 2006; Martin et al., 2011; Aldworth and Stopfer, 2015). Our goal is to identify the functional significance of certain characteristic neural response patterns observed as animals encounter sensory stimuli.

We are specifically motivated by experimental findings illustrating a rich taxonomy of stimulus-evoked, time-varying responses or sensory trajectories associated with both stimulus onset and offset (Stopfer et al., 2003; Bathellier et al., 2008; Saha et al., 2013b, 2017; Kay and Stopfer, 2006). It has been reported that the most noticeable change in activity of sensory neurons occur immediately following stimulus onset and then following stimulus offset (Mazor and Laurent, 2005; Stopfer et al., 2003; Raman et al., 2010; Bathellier et al., 2008) and these responses are in fact orthogonal to each other at a population level (Saha et al., 2017). In this work we are not attempting to ‘pattern-match’ these responses *per se*, but rather provide a functional interpretation of what these prototypical spatiotemporal response motifs achieve in terms of their ability to mediate stimulus detection. We address both through theory and validation the following: (i) Are there particular decoding objectives that explain the manifestation of observed sensory response motifs?; (ii) Are there physiologically plausible neural circuit architectures that are capable of realizing motifs generated by optimization of the objective in (i)?

The central theoretical premise of our paper is that activity of the early sensory network drives a latent representation of accrued evidence regarding presence or absence of stimulus. The neural responses thus encode not only detection, but also ‘undetection’ or withdrawal of the source of excitation. Adapting from approaches used in behavioral neuroscience, we formulate a dynamical decoder that uses high-dimensional, combinatorial input to generate a low dimensional latent representation. Thereafter, we use a top-down, normative approach wherein we generate the response motifs by minimizing an objective function. We formulate this objective function so that it emphasizes that latent representations should quickly and accurately convey information about peripheral stimulus in an energy efficient manner. Neural response motifs borne out of such an optimization problem bear morphological similarity with those observed *in vivo*, leading us to assert our claim of ascribed functional relevance.

Our work builds on a rich theory of olfactory coding focused on the question of how reliable and informative representations propagate through the sensory hierarchy (Zhang and Sharpee, 2016; Bhandawat et al., 2007; Schaefer and Margrie, 2007). We note especially, recent efforts to ascribe particular functions to the circuit architectures, including enabling stimulus reconstruction (Qin et al., 2019), categorization (Dasgupta et al., 2017) and novelty detection (Dasgupta et al., 2018). However, these findings have mostly been pursued in a static input-output domain, i.e., with tacit assumption of instantaneous, algebraic signal transformation. In contrast, we work in the space of sensory dynamics and time-varying representations, a topic that has received much attention from a descriptive standpoint (Laurent, 1996; Laurent et al., 2001; Rabinovich et al., 2000) but less so from the perspective of normative synthesis. In particular, we investigate through formal mathematical arguments what advantages the big switch between onset and offset responses achieve in the biological world, where organisms encounter stimuli in a dynamic fashion.

Our results show that our normative model produces emergent phenomena that predict many nuanced features of actual sensory network activity as observed in locust (Saha et al., 2017; Nizampatnam et al., 2018; Mazor and Laurent, 2005). Further, our normative synthesis procedure yields a set of network dynamics that is highly compatible with the known physiology of these circuit *in vivo*. We hereafter proceed to formulate our theoretical setup before presenting our key synthesis and validation results.

## Materials and Methods

### Computational model

#### Decoder and latent space

Our problem setup is premised on the canonical architecture of the insect early olfactory system (Kay and Stopfer, 2006; Masse et al., 2009), wherein chemical cues are transduced to neural signals through olfactory receptor neurons (ORNs) which then propogate activity via glomeruli to Projection Neurons (PNs) en route to higher brain areas. Our focus is on the dynamical transformations mediated by PNs and their local circuitry.

Our model for sensory tracking hinges on the definition of a latent space, ***ν(t)*** ≡ [*ν*_1_(*t*), *ν*_2_(*t*), …, *ν_m_*(*t*)] that contains information about stimulus identity. Each dimension of the latent space indicates accrued evidence regarding the presence of high-level stimulus features (Raman et al., 2011). The assumption is that such features provide an actionable representation of the stimulus that can then be used to enable higher-level processing and behavior. Specifically, PN activity, **x** ∈ ℝ^*n*^, is linearly decoded into the latent space via (1):

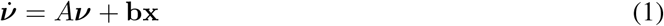

where *A* ∈ ℝ^*m×m*^ such that *σ*(*A*) ∈ ℝ^−^ and *σ*(*A*) represents the spectrum of *A*. Here, *A* = *−a***I**, where *a* > 0 and **I** ∈ ℝ^*m×m*^ is the identity matrix. The matrix **b** ∈ ℝ^*m×n*^ linearly mixes the contribution of PNs onto each dimension of the latent vector. Equation (1) imposes that the change in accrued evidence at any time *t* is proportional to the difference between the information encoded by representative PNs and evidence lost due to intrinsic leaky dynamics of the decoder. Figure 1A illustrates our problem setup, noting that we are focused on activity downstream of the ORNs (i.e., the dashed vertical line). Afferent activity from the receptor neurons is denoted by **r**(*t*). It is important to note that our approach does not assume any network structure nor dynamics for the PNs; rather, our goal is to synthesize these dynamics.

**Figure 1:**
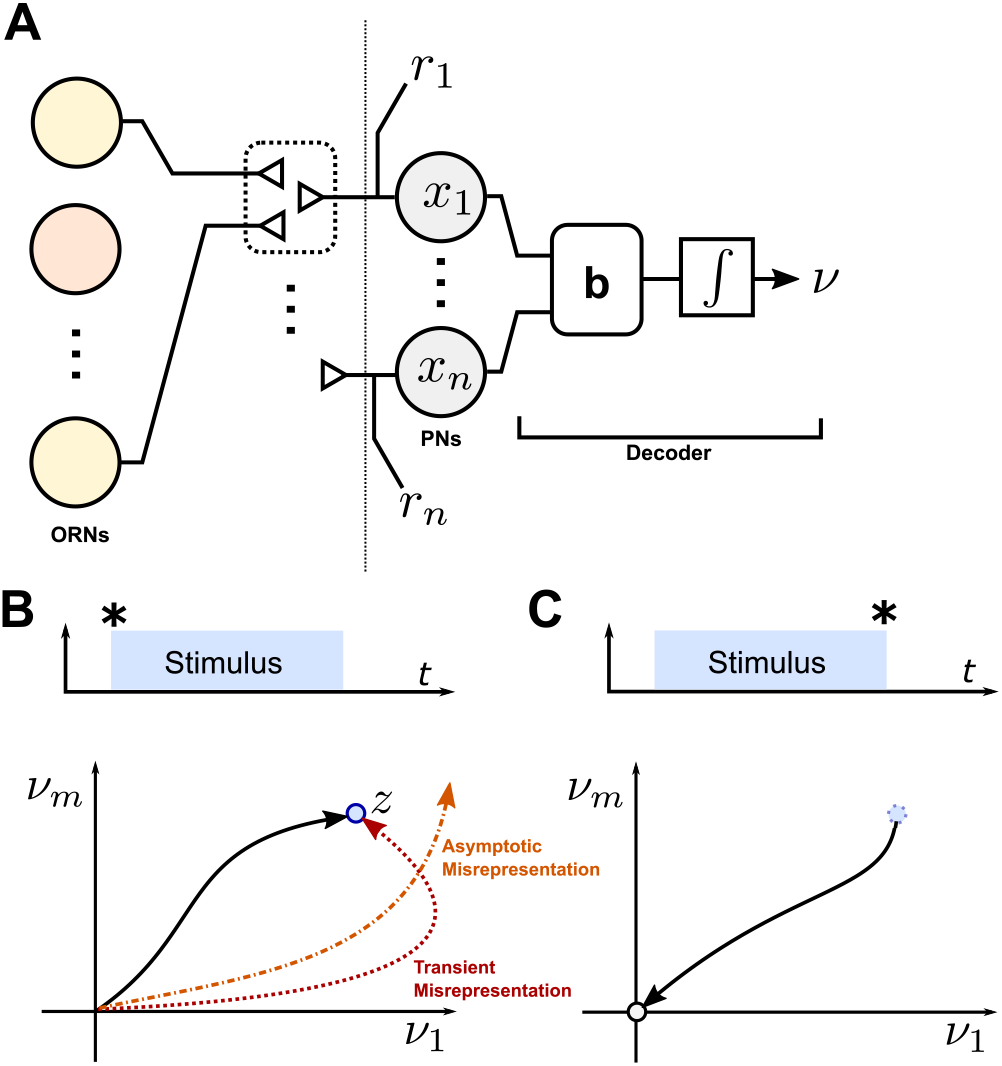
**A.** Schematic of signal flow through the early olfactory system. We consider response dynamics for the PNs subject to input **r**, which abstracts afferent signals from ORNs. **B**. We posit a decoding objective. At stimulus onset, the goal of PNs is to accurately drive a latent representation to a nominal target. **C.** At stimulus offset, the latent representation should return to neutral.

#### Enabling labile, accurate latent representations via sensory tracking

In particular, the formulation (1) allows us to introduce our normative premise, i.e., that the PN activity **x** drives accurate representations of accrued evidence, thus conveying information about stimulus presence and identity. Further, the dynamics of **x** should allow for fast transitions in ***ν***.

This idea is readily captured through the notion of tracking in the latent space. From the initial state ***ν*** = 0, we seek to construct dynamics for the PNs **x** such that ***ν*** can quickly and accurately reach an arbitrary point in the latent space, ***z***. Mathematically, we formulate a quadratic objective function

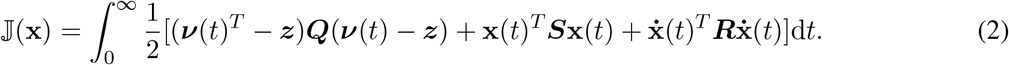

The terms in this objective function are highly interpretable from a biological perspective. The first term in the integrand penalizes the deviation of the decoded state from ***z***. The second term penalizes the energy of the PN response, while the last term penalizes large fluctuations in the PN response. The objective is formulated over an infinite horizon, i.e., there is no bias or expectation regarding the amount of time that the representation is to be tracked.

The objective and normative optimization is schematized in Figures 1B,C. At the stimulus onset (Figure 1B) the latent state is to track some nominal representation ***z***. On termination of the stimulus the latent state continues tracking ***z*** which is now positioned at the origin of the *m*-dimensional latent space signifying absence of a stimulus. Here, we considered a pulsatile stimulus structure in accordance with the natural olfactory environment, where an organism perceives olfactory stimulation as pockets of odor plumes (Vickers, 2000). Furthermore, in this work our focus was on the neural dynamics downstream of the receptor neurons (see Fig. 1 A), therefore, we assumed that each stimulus is presented at a fixed intensity. The matrix ***Q*** in our setup enforces that the tracking in the latent space occur accurately, without transient or asymptotic misreprsentation (see Figure 1B). The matrix ***S*** imposes penalty on the energy expenditure while the matrix ***R*** regularizes rapid fluctuation in firing rate activity. Upon withdrawal of the stimulus, (Figure 1C) the representation returns to neutral. In essense, this model embeds the idea of active stimulus detection and ‘undetection’. The question at hand is how **x** should be specified (according to 2) for robust and accurate detection of odor stimuli.

Table 1 summarizes all mathematical symbols used in this paper as well as their interpretation.

**Table 1:** Table of math symbols and their interpretation.

| Mathematical symbol | Meaning in the model | Significance in the olfactory system |
| --- | --- | --- |
| $\mathbf{r}(t)$ | Afferent input to PNs | Activity of olfactory receptor neurons (ORNs) |
| $\mathbf{x}(t), \mathbf{x}_o(t)$ | Activity of neural units driving the latent decoder | Firing rate of Projection Neurons(PNs) with baseline activity set to zero. |
| $\mathbf{x}_i(t)$ | Activity of auxiliary population of neurons | Firing rate of Local Neurons (LNs) with baseline activity set to zero |
| $\nu(t)$ | Decoded latent space activity: each dimension indicates accrued evidence regarding presence of high-level stimulus features | Intermediate representation of stimulus information that drives downstream behavioral/motor responses |
| $A, \mathbf{b}$ | Dynamics of the latent space decoder. $A$ represents the internal dynamics of the decoder. Weights of matrix $\mathbf{b}$ represent combinatorial encoding | |
| $z$ | A fixed representation associated with a particular stimulus | |
| $Q$ | Penalty on accuracy of latent representation | |
| $S$ | Penalty on neural resources i.e., energy used | |
| $R$ | Regularization on rapid fluctuation in firing rate activity | |
| $W_s$ | Connections mapping slow processing of $\mathbf{x}(t)$ onto its dynamics | |
| $W_f$ | Connections mapping fast processing of $\mathbf{x}(t)$ onto its dynamics | |
| $W_o^i$ | Excitatory connections from the representative PNs to the auxiliary population | Excitatory synaptic connections from PNs to LNs |
| $W_i^o$ | Inhibitory connections from the auxiliary population to the putative PNs | Inhibitory synaptic connections from LNs to PNs |
| $W_i^i, W_o^o$ | Interpopulation interactions | Synaptic connections among PNs and LNs. |
| $\overline{\mathbf{W}}, \hat{\mathbf{W}}$ | Computational parameters (matrix) that must satisfy the following: $\hat{\mathbf{W}}\overline{\mathbf{W}} = \mathbb{I}$, $vec(\overline{\mathbf{W}}) \geq \mathbf{0}$ and $vec(\mathbf{W}_s\hat{\mathbf{W}}) \leq 0$ | |
| $C$ | Fraction of neurons responding to both red/blue stimuli | Percentage of Projection neurons that respond to more than one sensory stimulus |
| $\eta(t)$ | Unit variance Gaussian noise for the network model developed through Stochastic LQR formulation | Background noise in neural response |
| $t_s$ | Duration of stimulus period | Duration of stimulus period |

### Experimental Design

#### Odor stimulation

We followed the similar protocol as in our previous studies for odor stimulation (Saha et al., 2013b, 2017). The odor solutions were diluted in mineral oil (from Sigma-Aldrich) to achieve 1% dilution (v/v). 20 ml of diluted odor solution was placed in a 60 ml sealed glass bottle with inlet and outlet lines. A constant volume (0.1 L min^−1^) from the odor bottle headspace was injected into the carrier stream using a pneumatic pico-pump (WPI Inc., PV-820) during odor presentations. A vacuum funnel was placed right behind the animal preparation to ensure the removal of odor vapors. Odor presentations were 4 s long. Each trial was 40 s in duration with an inter-trial interval was set to 20 s.

#### Extracellular recordings

Post fifth instar locusts (*Schistocerca americana*) of either sex were selected and were immobilized first, and then the brain was exposed, desheathed as reported in previous studies (Saha et al., 2013a; Laurent and Davidowitz, 1994; Brown et al., 2005). Extracellular recordings of PNs were performed using NeuroNexus 16-channel, 4×4 silicon probes. Impedances of all the electrodes were kept in the 200-300 kΩ range. Raw extracellular signals were amplified at 10k gain using a custom make 16-channel amplifier (Biology Electronics Shop; Caltech, Pasadena, CA), filtered between 300 Hz and 6 kHz, and acquired at 15 kHz sampling rate using a custom-written Labview software.

#### Intracellular recordings

Animal preparation for intracellular recordings was same as that used for extracellular recordings. Patchclamp (in vivo) recordings from PNs were performed using the pipettes filled with locust intracellular solution (Laurent et al., 1993): 155mM K aspartate, 1.5mM MgCl_2_, 1mM CaCl_2_, 10mM HEPES, 10mM EGTA, 2mM ATP disodium salt, 3mM D-Glucose, 0.1mM cAMP. All these chemicals were purchased from Sigma-Adrich. pH of the patch solution was adjusted to 7.0 using 1M NaOH and Osmolarity was adjusted to 320-325mM range using Sucrose. The impedances of electrodes ranged between 5–15 MΩ. Raw voltage traces were amplified (Axoclamp 900a) and acquired at 16 kHz sampling rate using a Labview software.

## Results

### Sensory tracking is achieved by a biologically plausible network architecture

The objective function (as in Eq. (2)) taken together with the decoder dynamics (Eq. (1)) results in the following optimization problem.

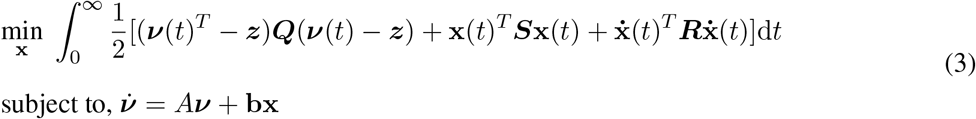

The solution of this optimization ((3)) yields the following PN dynamics:

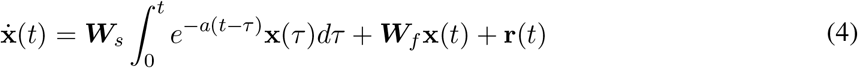

Eq. (4) reveals the existence of a network that incorporates both slow and fast processing (via ***W***_*s*_ and ***W***_*f*_, respectively) in response to an external stimulus. Also present is dependence on the exogenous stimulus via **r**(*t*), the readout from the first-level receptor neurons. However, how the slower timescale is achieved *in vivo* is not explicitly explained by (4). The question here is: can a biologically plausible first order network architecture be realized by decomposition of (4) into first order rate equations? We find that by introducing an auxiliary population of neurons which share reciprocal connections with the second order PNs, this can be achieved. Eq. (5) below is a representative network architecture arising from (4) that produces the optimal motif as seen in Figure 3.

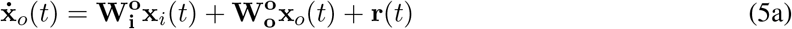

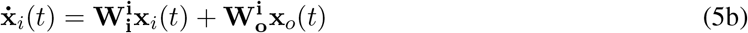

**Figure 2:**
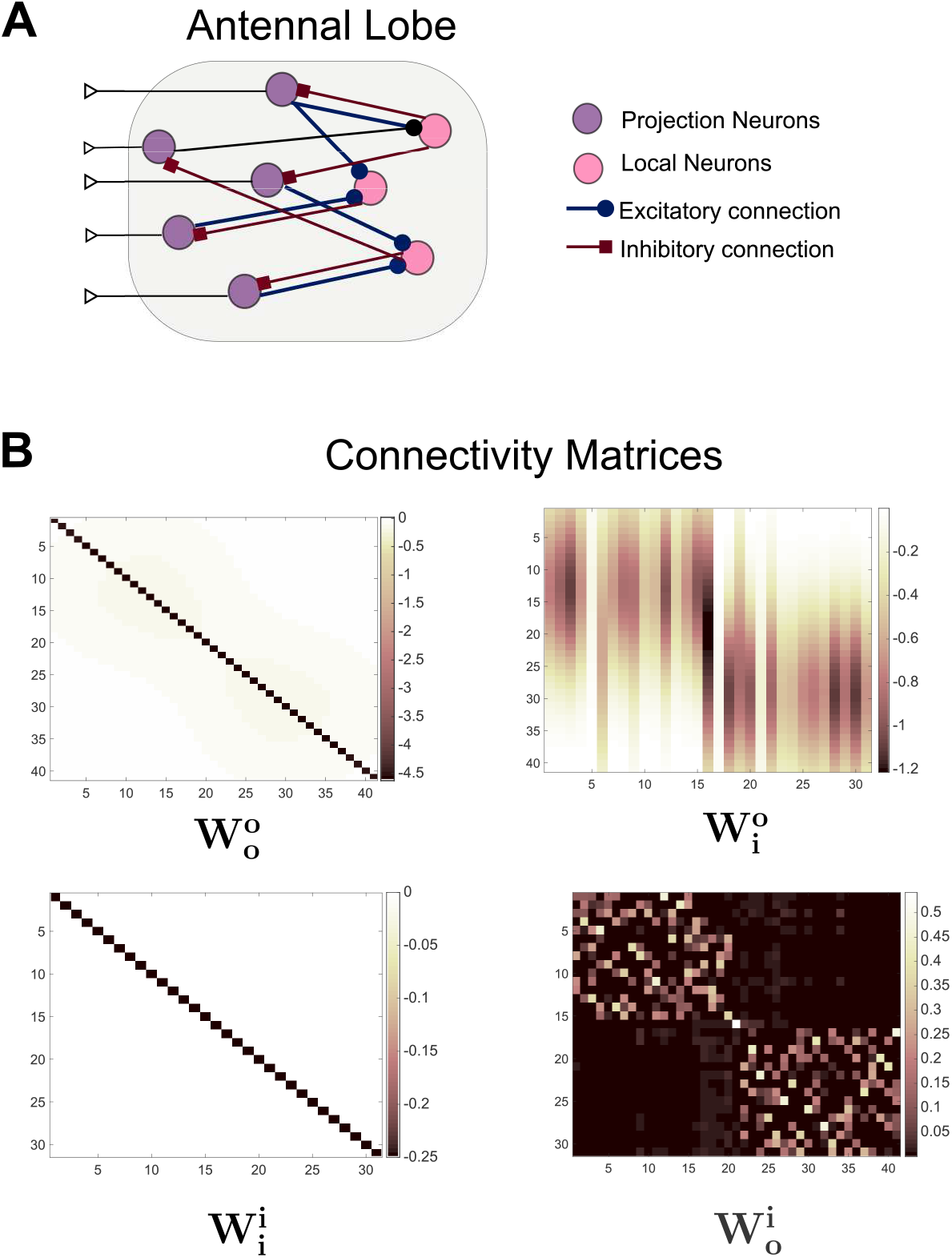
**A.** Schematic of synaptic interactions between PNs and LNs in the Antennal Lobe of insects. The PNs provide excitation to LNs, whereas LNs impose inhibitory control on PNs. **B.** Connectivity matrices arising for a particular choice of 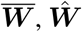 in our mathematical model shows the synaptic interactions between the two populations. It is consistent with the canonical excitatory-inhibitory scheme observed in sensory networks *in vivo*.

**Figure 3:**
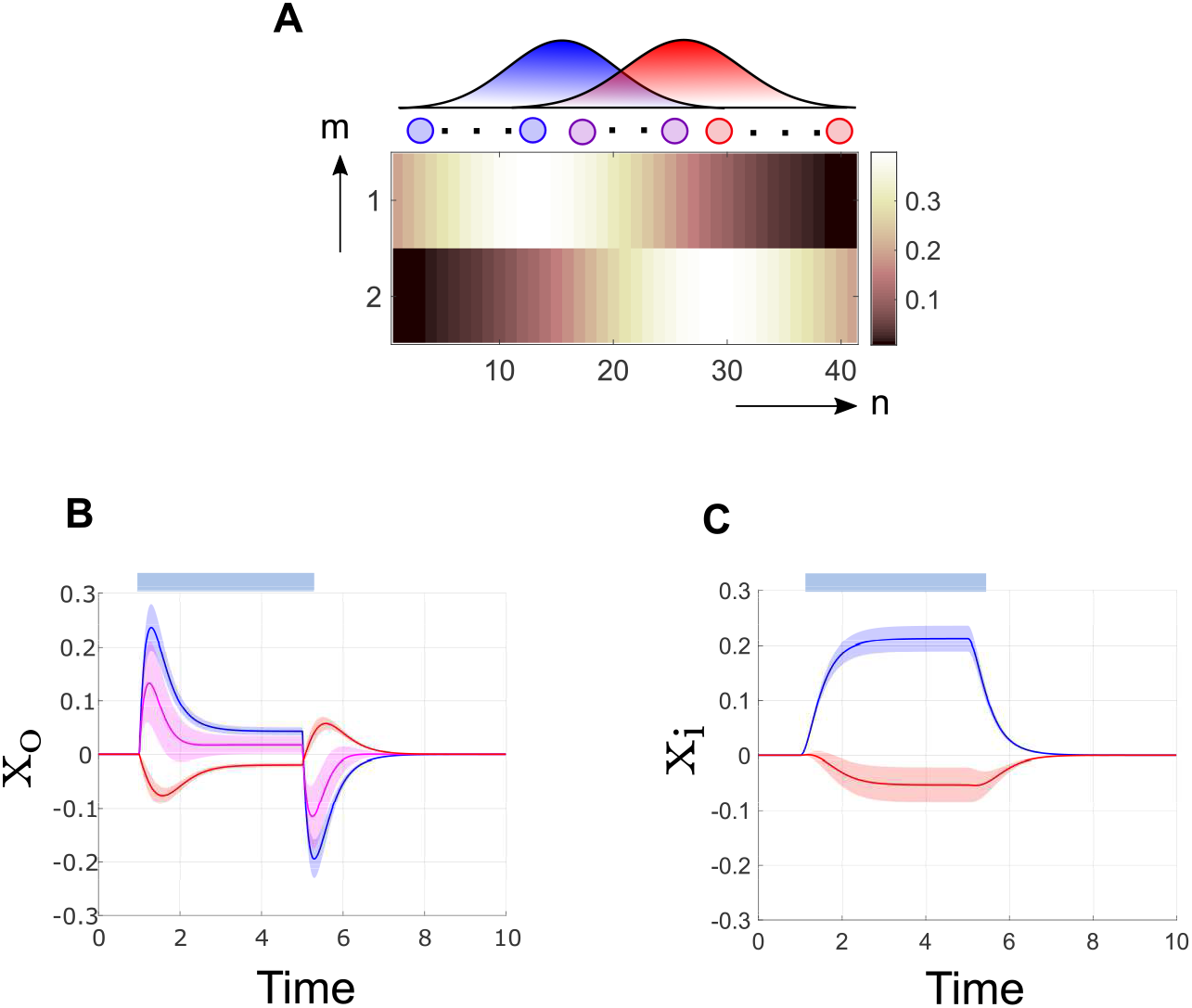
**A.** Combinatorial encoding is represented through the weights of matrix **b** in our model. Each row determines the contribution of the *n* = 41 neural units to evidence accrued corresponding to the high-level feature. **B-C** Response motifs produced by the two populations (PNs and LNs) in presence of a blue pulse. Depending on their tuning to blue/red stimulus, the neurons may be excited or inhibited respectively. **B.** PN activity displays two phasic transients one on stimulus onset and the other on stimulus withdrawal, in between these transients the activity is steady but reduced in amplitude. **C.** LN activity is present chiefly as long as the stimulus is in action.

Here, 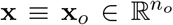 is the PN activity and 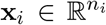 is the activity of the auxiliary population, and *n_o_* and *n_i_* are the number of neurons belonging to each population subgroup. The connectivity matrices in (5) are analytically specified as 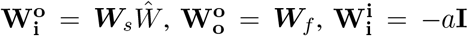 and 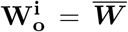, where 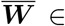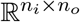 and 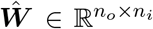. In the insect Antennal Lobe(AL), PNs extend excitatory connections onto an auxiliary population of neurons known as Local Neurons(LNs) which in turn provide inhibitory control on the PNs (Shang et al., 2007; Kay and Stopfer, 2006; Wilson and Laurent, 2005). In our network architecture, connections between PNs and the auxiliary population are encapsulated in the matrices 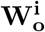 and 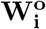. The structure of these matrices depend upon choice of 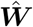 and 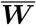 (chosen such that 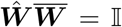). We set the dimensions of these matrices such that *n_i_* < *n_o_* as cellular studies have reported that typically the size of the population of LNs is smaller than that of PNs (Laurent, 1996). 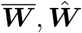 are chosen through a constrained iterative optimization scheme (see Appendix for details). The constraints for this secondary optimization problem are stated on the basis of the canonical excitatory-inhibitory structure found in the insect antennal lobe: (i) 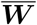 (PN to LN) must be non-negative and (ii) 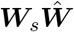 (LN to PN) must be non-positive. We initialized our proposed algorithm by selecting 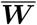 to be a positive random block matrix such that its pseudoinverse existed and we set the initial choice for 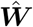 as 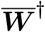, the Moore Penrose pseudo-inverse of 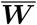. The choice of such an initialization was fashioned after our prior knowledge regarding existence of stimulus specific tuning of the PNs (Saha et al., 2017; Assisi et al., 2011). It is important to emphasize here that the solution of this iterative scheme is non-unique due to the structure of the problem (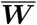 is a wide matrix and 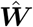 is its left inverse). In other words, there will in general be many network architectures that realize the optimal motif from (3). The common trend observed across various solutions (each a mathematically valid network architecture) is that the connections between PNs and LNs are essentially random (salt and pepper). This is consistent with *in vivo* observations wherein each PN extends its efferent terminals onto only a subset of the auxiliary population and receives inhibitory control only from a subset of the population (Carey and Carlson, 2011; Bazhenov et al., 2001; Saha et al., 2017).

Inter-population interactions of PNs and LNs are captured in the matrices 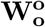 and 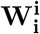. The diagonal terms in these matrices represent the self-decay dynamics associated with each neuron. Our construction also revealed very weak inhibitory connections within the PN population pool. Such connections are not known to exist experimentally, and we surmised that these small negative weights could be proxies of fast inhibitory synapses (Bazhenov et al., 2005, 2001) occurring elsewhere in the antennal lobe (see also Discussion).

### Network dynamics produce stimulus onset and offset responses that are observed *in vivo*

While the architecture of the normative model is consistent with that of the antennal lobe, it is not yet clear whether its dynamics and activity also matches the types of responses observed *in vivo*. To study this issue, we set up a illustrative model for binary discrimination in a combinatorial setting. We emphasize here that the choice of a two-dimensional state space is made without loss of generality (see Appendix for an example of a system with higher dimensional latent space).

The combinatorial nature of the encoding space is embedded in **b** (see Eq. (1)), which determines the tuning of individual neurons. In the example network, we used Gaussian tuning curves as per Figure 3, so that some neurons respond preferentially to one stimulus or the other (here, visualized as red vs. blue), while some do not exhibit a tuning preference (referred to as untuned). It is observed that the PNs generate two significant phasic bursts of activity - one following odor onset and the other on odor termination (see Figure 3B). Moreover, the on and off responses have orthogonal orientations in a dimensionality-reduced space and are negatively correlated (see Figure 5B, C and Appendix for dimensionality reduction details). While spatio-temporal patterns of stimulus evoked activity in the Antennal Lobe(AL) of insects has been studied previously in literature (Laurent et al., 1996; Raman et al., 2010; Saha et al., 2017), our normative formulation provides an insight into the functional relevance of such response patterns, i.e., these motifs allow for robust and accurate stimulus representations for downstream processing in an energy-efficient manner.

Depending upon their tuning, PN responses fall into one of the following categories: (i) a rapid increase in firing at the onset of stimulus that subsequently settles to a steady state response of reduced amplitude; on removal of stimulus there exists a brief period of inhibited activity (below baseline) before the system returns to baseline, or, (ii) a state of suppressed firing activity through the entire duration of stimulus, but excitatory (above baseline) firing activity when the stimulus is withdrawn (see Figure 3B, C). On the other hand, the auxiliary neurons (i.e., the putative LNs) display tonic activity in response to stimuli, returning to baseline gradually when the stimulus is no longer active. While the tuning of PNs has been studied extensively (Wilson et al., 2004; Stopfer et al., 2003), whether similar combinatorial architecture exists in the LNs is yet to be fully characterized. In Figure 3C, the blue/red coloration does not imply prior tuning, but simply indicated whether the neurons were excited/inhibited by the stimulus (Saha et al., 2017).

Interestingly, these predicted response dynamics have been observed in recordings from the locust olfactory system, wherein recent findings suggest both phasic and tonic temporal responses associated with stimulus onset and offset (Saha et al., 2017). Specifically, experimental observations indicate that when a pulse of stimulus is provided, the neurons in the antennal lobe exhibit a phasic transient followed by a tonic activity for the duration of the stimulus (referred to as *On response*). On termination of the pulse, there is another short-lived burst of activity before activity slowly returns to the quiescent regime (an *Off response*) (Saha et al., 2017). Figure 4 shows intracellular voltage traces recorded from four representative projection neurons (PNs) in the locust antennal lobe. As can be noted, the stimulus-evoked responses in these PNs occur either when the stimulus is present followed by hyperpolarization (ON-type), or following the termination of the stimulus (OFF-type; note that membrane potential is hyperpolarized when the odor was presented). This segregation of stimulus-evoked PN responses into these distinct categories is maintained even when a larger number of neurons probed with a wider panel was considered (Saha et al., 2017). It should be mentioned here that the ensemble response vectors during onset and offset responses have distinct geometric orientations and are negatively correlated with each other as reported by both model simulations and experimental recordings (see Fig. 5A-C).

**Figure 4:**
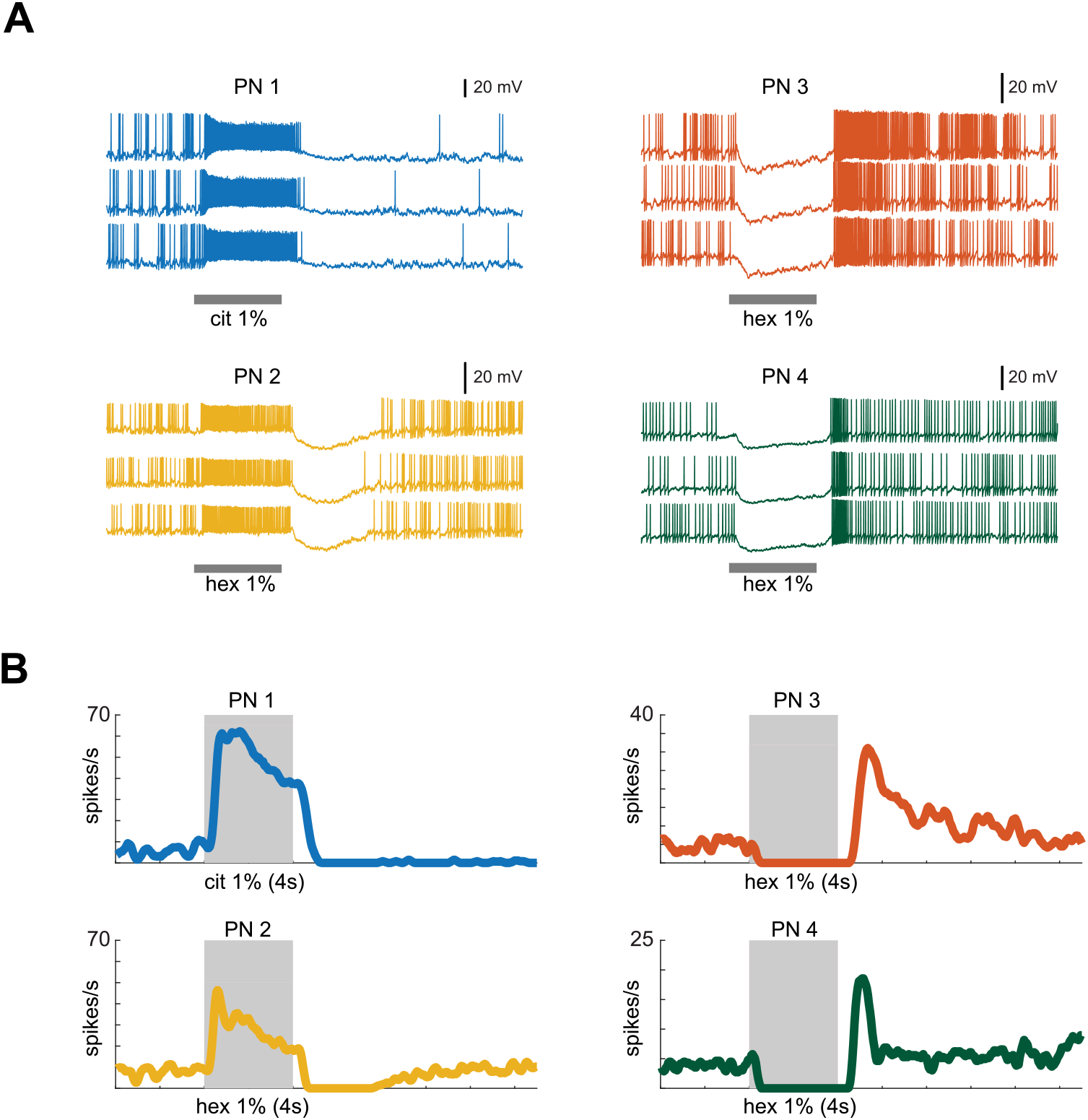
**A.** Intracellular voltage traces of four different projection neurons (PNs) are shown (different colors). Shaded gray box in each panel shows the 4 s duration of odor exposure (cit – citral, hex – hexanol; both delivered at 1% v/v dilutions) and the black line represents the scale bar (20 mV). Three traces in each panel correspond to three consecutive trials of recording, revealing that the temporal patterns are consistent between trials. PN1 and PN2 responded during the presentation of odor (*On response*), and PN3 and PN4 responded after the termination of odor (*Off response*). **B.** PN firing rates in 50 ms time bins were averaged across trials and are shown in **A**.

**Figure 5:**
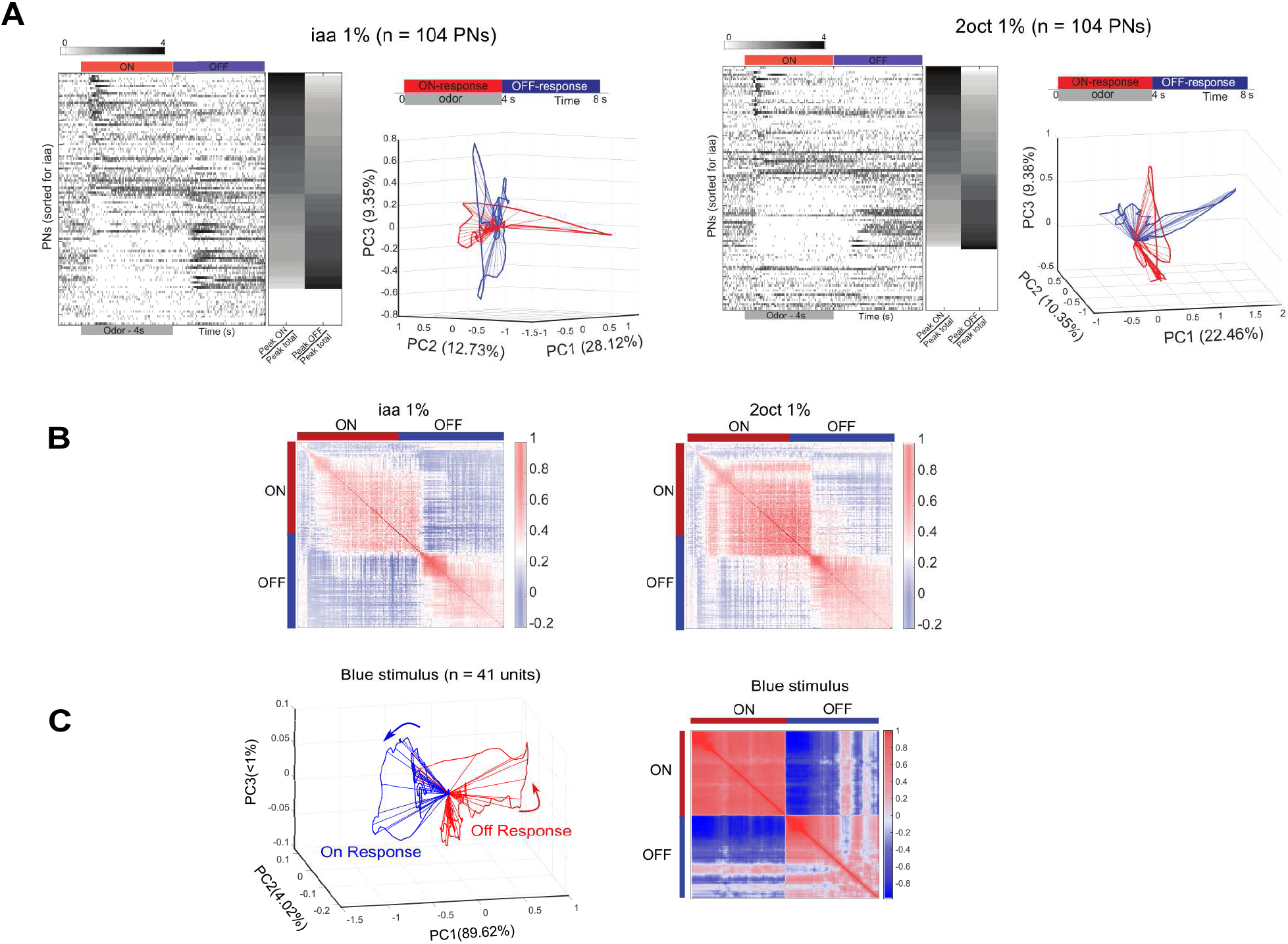
**A.** Mean PN firing rates (50 ms time bins and averaged across 10 trials) are shown for each PN (single row), and across the entire PN ensemble (different rows). The 4 s when the odorant was present (i.e. On period) and 4s after the termination of the stimulus (i.e. Off period) is identified using a red and a blue bar respectively. PNs are ordered based on the difference between the peak firing rates during ON and OFF epochs (Saha et al., 2017). Non-responsive neurons are shown at the bottom. Normalized peak firing responses during ON and OFF periods for each PN are shown on the right of the panel as a color bar. Neural ensemble response trajectories after PCA dimensionality reduction are shown. Red trajectory corresponds to the 4 s when iaa was presented, and the blue trajectory corresponds to the 4 s after termination of the stimulus. Similar plots for two different odorants (Left)isoamyl acetate at 1% v/v and (Right)2-octanol at 1% v/v are shown. **B.** Correlation between ensemble response evoked by an odorant; the ON and OFF responses are negatively correlated. **C.** PCA trajectories and correlation maps generated during 4s stimulus(blue) onset and 4s following stimulus withdrawal generated by the noisy model are shown.

### Orthogonalization of population response achieves robust latent representation

In the previous section we have explored how the PNs achieve stimulus detection in a latent space via spatio-temporal response patterns. We sought to understand how PN response patterns that are optimized for stimulus detection affect other aspects of sensory processing. One prediction that was made by the model pertains to processing stimuli that are encountered in sequence, often without sufficient inter-stimulus interval for the system to recover fully. Our model revealed that in such situations resultant PN responses emphasize albeit more strongly on distinct orientations for distinct stimuli (in the reduced dimension space) strongly (see Fig. 6 A-C). This finding has been previously referred to as the contrast enhancement computation (Nizampatnam et al., 2018) that highlights the uniqueness of the current stimulus with respect to prior distractions (see Fig. 6E). Here, we used our defined latent space of intermediate representation to analyze the need for such patterning. It turns out that what appeared as contrast enhancement/ novelty detection through activity of PNs in fact enabled maintenance of robust latent representation (see Fig. 6D) in the face of a variety of background distractions.

**Figure 6:**
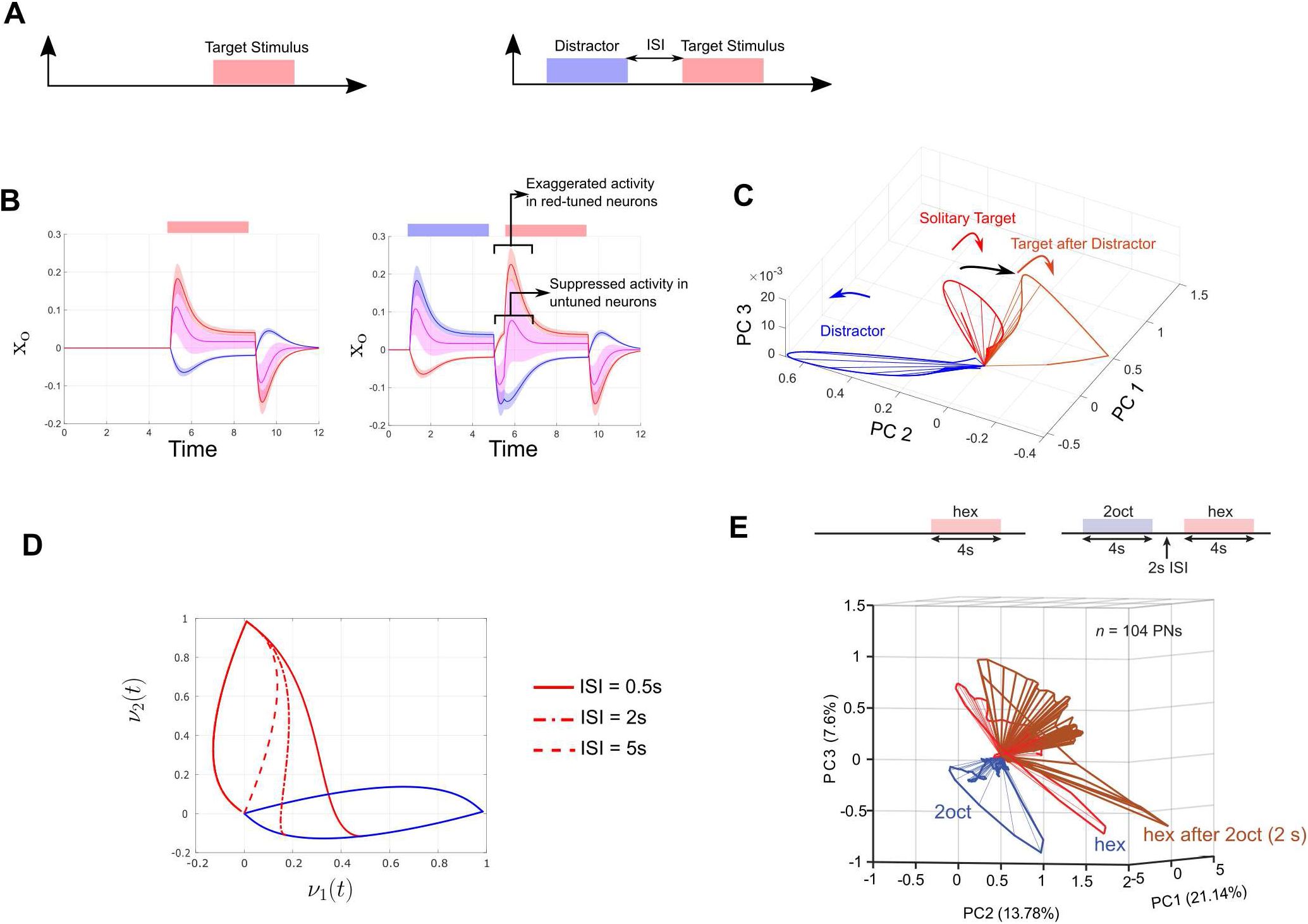
**A.** The synthesized model is excited by (a) a solitary red target pulse or (b) a blue distractor pulse followed by a red target pulse. **B.**Change in neural activity due to presence of distractor cue. **C.** PN activity visualized after dimensional reduction: response trajectory in the distractor-target sequence implements contrast enhancement (Nizampatnam et al., 2018). **D.** Latent state representation for a range of Interstimulus Interval. The model predicts that robust latent realizations are possible even when interstimulus intervals are very small due to contrast enhancement computations. **E.** Similar plots as in Fig. 5A are shown for a sequential presentation of hexanol (hexanol after 2s ISI shown as brown trajectory). Neural response trajectories for solitary presentations of 2-octanol (blue trajectory) and hexanol (red trajectory) are also plotted for comparison. Note that compared to the red trajectory, brown trajectory is more distant from the blue trajectory.

Importantly, the objective function in our optimization framework (Eq. (2)) stressed only on formation of accurate latent representation in an energy efficient manner for a single stimulus at a time. Regardless, this framework predicted response features that are observed in the biological world where stimuli are often perceived under a variety of contexts (Nizampatnam et al., 2018). This functional interpretation of what is achieved by neural response modulation when stimuli appear in conjuction with a multitude of distractions is an interesting outcome of this study.

### Active reset enables consecutive fast and accurate representations

As suggested by the above finding, the synthesized network embodies the notion of *active-reset* via its off response. We further analyzed the functional advantage of such distinct reversal of firing trends at the conclusion of the active stimulus period by systematic simulations of our model. We considered here two model alternatives (a) the synthesized network and (b) a network whose activity ceases immediately on withdrawal of the stimulus and the decoder relies only on its natural decay to return to neutral (‘passive reset’). We simulated each of these models with a sequence of stimuli (see Fig. 6A right panel). It turned out that the active reset mechanism mediated via the off response prepares the system promptly for the next incoming excitation. In the absence of such swift reversal of firing activity at the conclusion of a stimulus the system ‘recovered’ at a much slower rate leading to confounding latent representation of succeeding stimuli (see Fig. 7A). Clearly, the extent of misrepresentation depends on the degree of passive reset (i.e., the parameter *a*) as well as the time allowed for the system to recover between pulses (see Fig. 7B; the definition of accuracy in this context is included in the Appendix).

**Figure 7:**
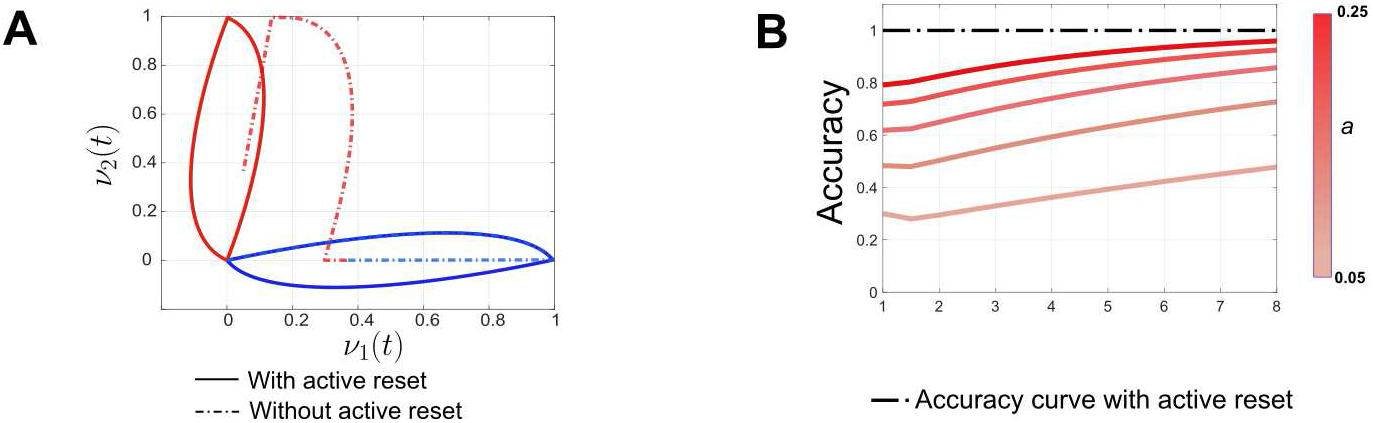
**A.** In absence of active reset, latent state trajectory evolves with a bias introduced by the previous cue. The amount of bias introduced in the system depends upon the intrinsic dynamics of the decoder controlled by *a* and the time between two stimuli. A comparison is made with a system implementing active reset (gray line).

**Figure 8:**
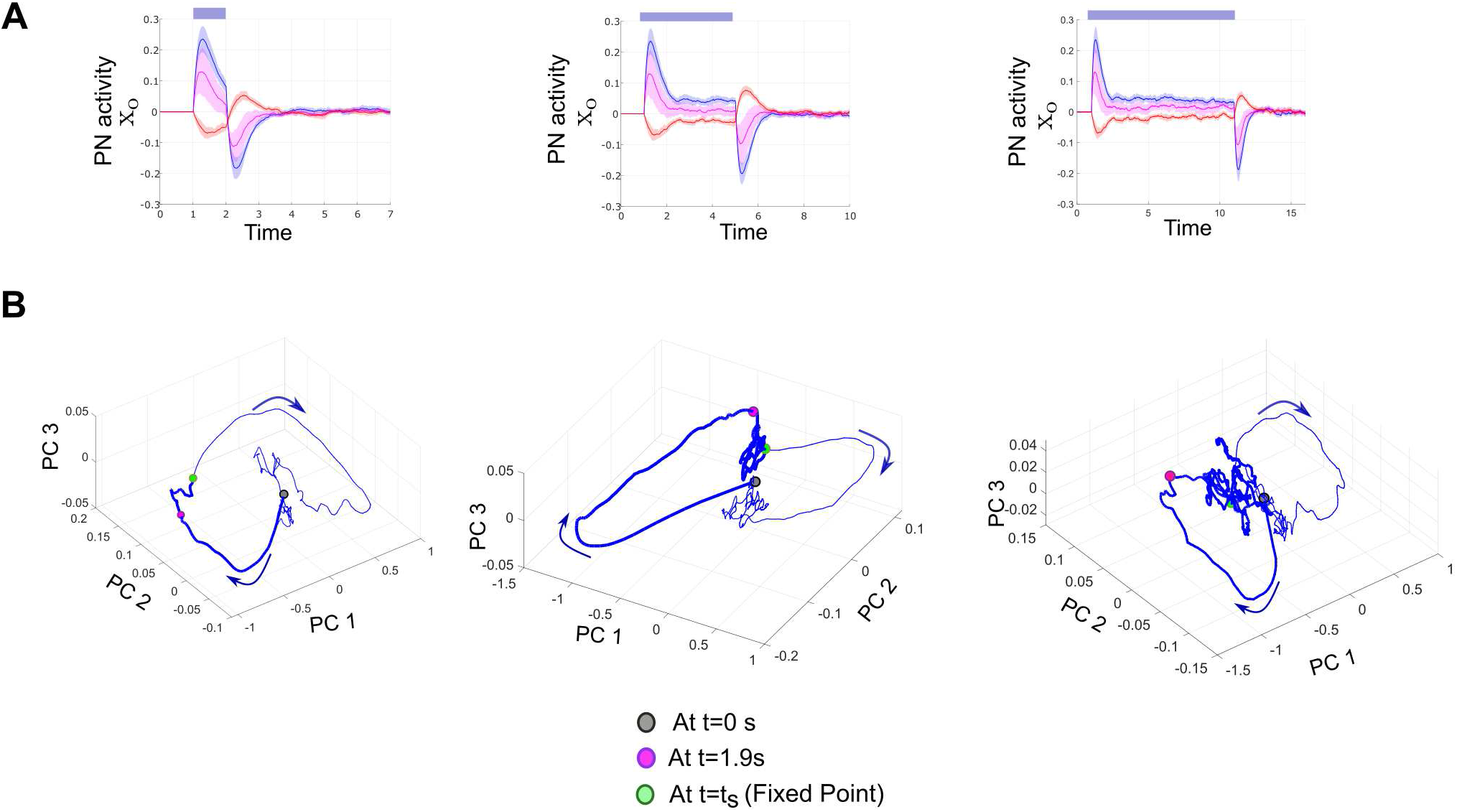
**A.** PN population response for different lengths of stimulation i.e., *t_s_* = 1s, 4s, 10s. **B.** PN trajectories as visualized in the PC space. Grey circle indicates the initial point, pink circle indicates the end of transient(phasic) response period and Green Circle indicates the fixed point to which the population response converges asymptotically in presence of a stimulus. The thicker part of the trajectory indicates the period for which stimulus was on. The arrows indicate the direction of traversal.

### The optimal response motif predicts the existence of a fixed point attractor

It has been found that PN population response comprising of phasic/tonic activity patterns when analyzed in the reduced Principal Component space trace distinct, odor specific trajectories. Starting from the origin or a neutral point, each odor-evoked trajectory quickly coalesced to a specific ‘fixed-point’ in the neural state space (Galán et al., 2004; Mazor and Laurent, 2005). The model in this study is designed to ‘track’ a nominal representation in the low dimensional decoded latent space with high confidence. In doing so, the derived optimal neural responses displayed a sharp transient activity at both stimulus onset and offset (see Fig. 3B); in the period between these transients the PNs responded to the stimulus by a steady state response of low amplitude. Therefore, the model indeed predicts the existence of an odor-specific fixed-point attractor in the neural state space. Beginning from a neutral state it took only a few milliseconds to be in the proximity of the attractor determined by stimulus identity; likewise, on withdrawal of the stimulus it is the transient activity that propeled the system to quickly return to its neutral regime. The interesting observation here is the speed with which the neural response mediated detection and ‘undetection’, as our optimization framework (2) is formulated to produce representations over an infinite time horizon without any bias towards when specific events (i.e., detection) should occur.

### Encoding performance degrades gracefully with respect to increasing tuning overlap

Combinatorial coding in olfaction is ubiquitously present (Kundu et al., 2016; Menini, 2009) as this scheme provides efficient utilization of limited neural resources for identification and processing of a very large number of odors. The question is, how robust is the performance of the model across different degrees of overlap in the tuning curves? In our mathematical setup, *C* is the fraction of neurons that responds to both red/blue stimuli with no overt preference to either stimuli; therefore it also indicates the extent of overlap between the tuning curves or similarity between the feature space of afferent stimuli. The synthesized model predicted that the network can maintain accurate representations of the stimuli for a large range of values of *C* (see 9A, B). Beyond this range of overlap however, there is a rapid decline in the quality of representation. It is evident that systems which endured large expenditures in terms of energy and maintained strict demands for error minimization were more tolerant to increased overlap in the combinatorial space (see Figure 9B) than their counterparts.

**Figure 9:**
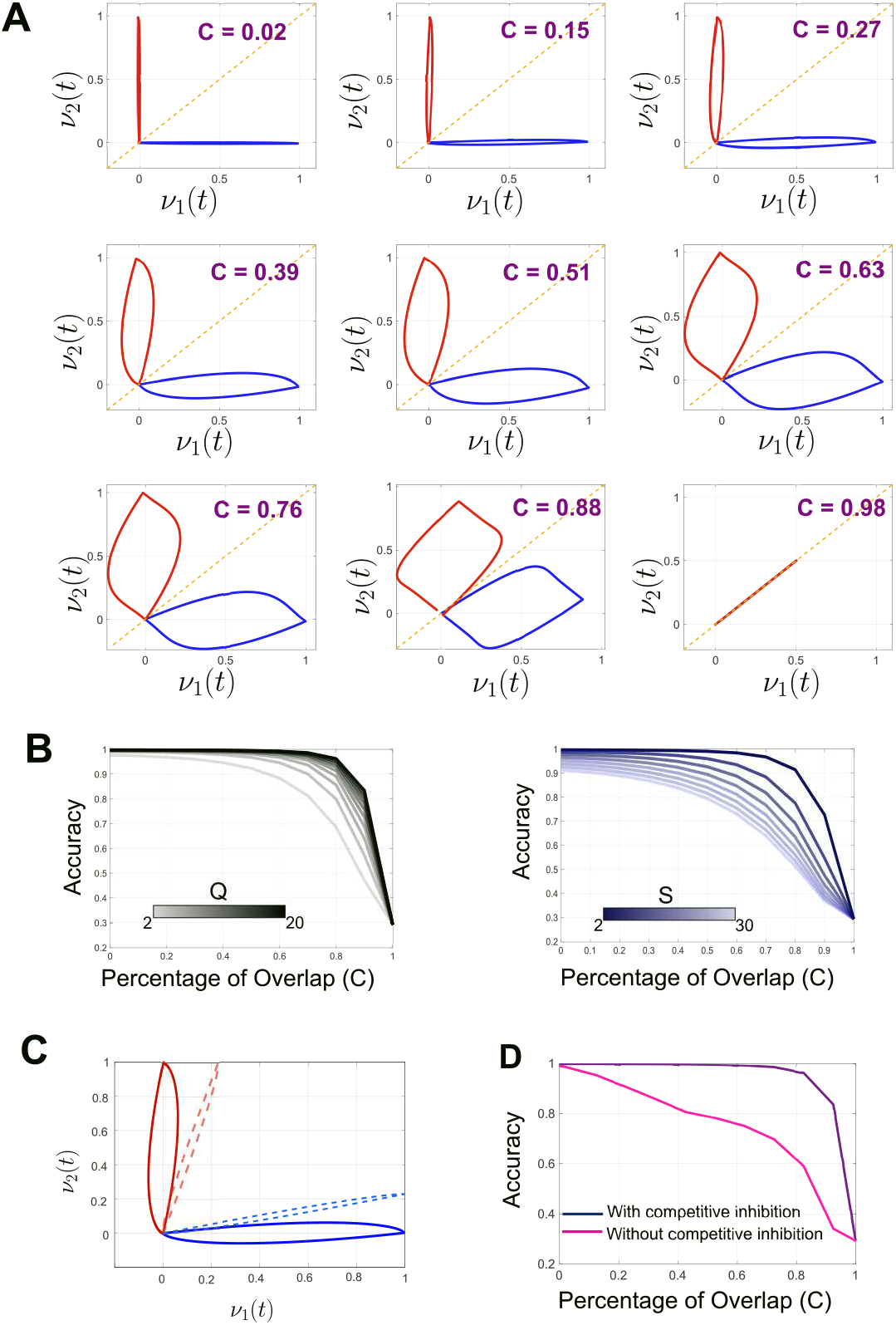
**A.** Trajectories in the latent space evolve with reduced spatial separation with increased overlap between Gaussian tuning curves embedded in **b**. However, the degree of misrepresentation (inversely related to spatial separation) is appreciably low across a wide range of overlap. **B.**Accuracy of the normative model as a function of the percentage of overlap. We notice that after a critical point, the performance degrades drastically. This point along the horizontal axis is a function of penalty matrix ***Q*** (left) and regulariztion matrix ***S*** (right). **C.** Competitive inhibition is pivotal for ensuring accurate representation. (left) In absence of competitive inhibition, the trajectories corresponding to red and blue stimulus are more proximal to each other (dashed line). **D.** Comparison between systems with and without competitive inhibition. 50% of the neurons responsive to the red stimulus are switched off while a blue stimulus was provided. The accuracy of the system with impaired competitive inhibition declined rapidly.

In this context the next question is, can this graceful deterioration in performance be attributed to some embedded feature of the optimal motif? On analysis we found that in fact it is the activity of the competing neurons that alleviated both transient and asymptotic misrepresentation over a considerable range of overlap in the tuning curves. We have previously identified that the neurons preferentially selective to a given stimulus remain in a state of suppressed activity in presence of a competing stimulus (3B). When this activity is eliminated by reducing the contribution of the red-tuned neurons to the latent decoder to baseline, we observed that the latent state trajectory tracked ***z*** poorly. Furthermore, the inherent property of graceful degradation of the synthesized network also disappeared in absence of competitive inhibition (see Figure 9C,D).

### Optimal motifs mediate a trade-off between latency, energy and accuracy

The choice of the error penalty matrix ***Q*** and energy regularization matrix ***S*** in (2) shaped the optimal neural responses and subsequently influencde the performance of sensory tracking in the latent space. Intuitively, as ***Q*** is scaled to higher values, stricter demands are made to ensure high fidelity tracking. Whereas, increasing the scale of ***S*** encourages greater degree of conservation of energy. The notion of a performance trade-off here is embedded therefore in the choice of matrices ***Q*** and ***S*** (see Figure 10A,B) as they share antipodal relationship with respect to accuracy as well as latency (time to reach 80% of the nominal evidence level). In the appendix of this paper we have reported how choosing a cosine metric to measure the quality of latent representation can be used. Figures 10C,D illustrates the response morphology associated with different penalties. This indicated that low penalty on tracking error and high regularization on energy produced smaller overshoots on stimulus onset. Such responses, while still capable of forming latent space representations, did so more slowly and less accurately(see cyan squares on surface plots of 10A,B). Phasic overshoot, conversely, in indicative of a network designed to promote accurate and fast representations (see pink circles on surface plots of 10A,B). Thus, our finding is compatible with the argument that short stimulus exposures are sufficient for olfactory discrimination (Uchida and Mainen, 2003), at the cost of higher energy expenditure within sensory networks.

**Figure 10:**
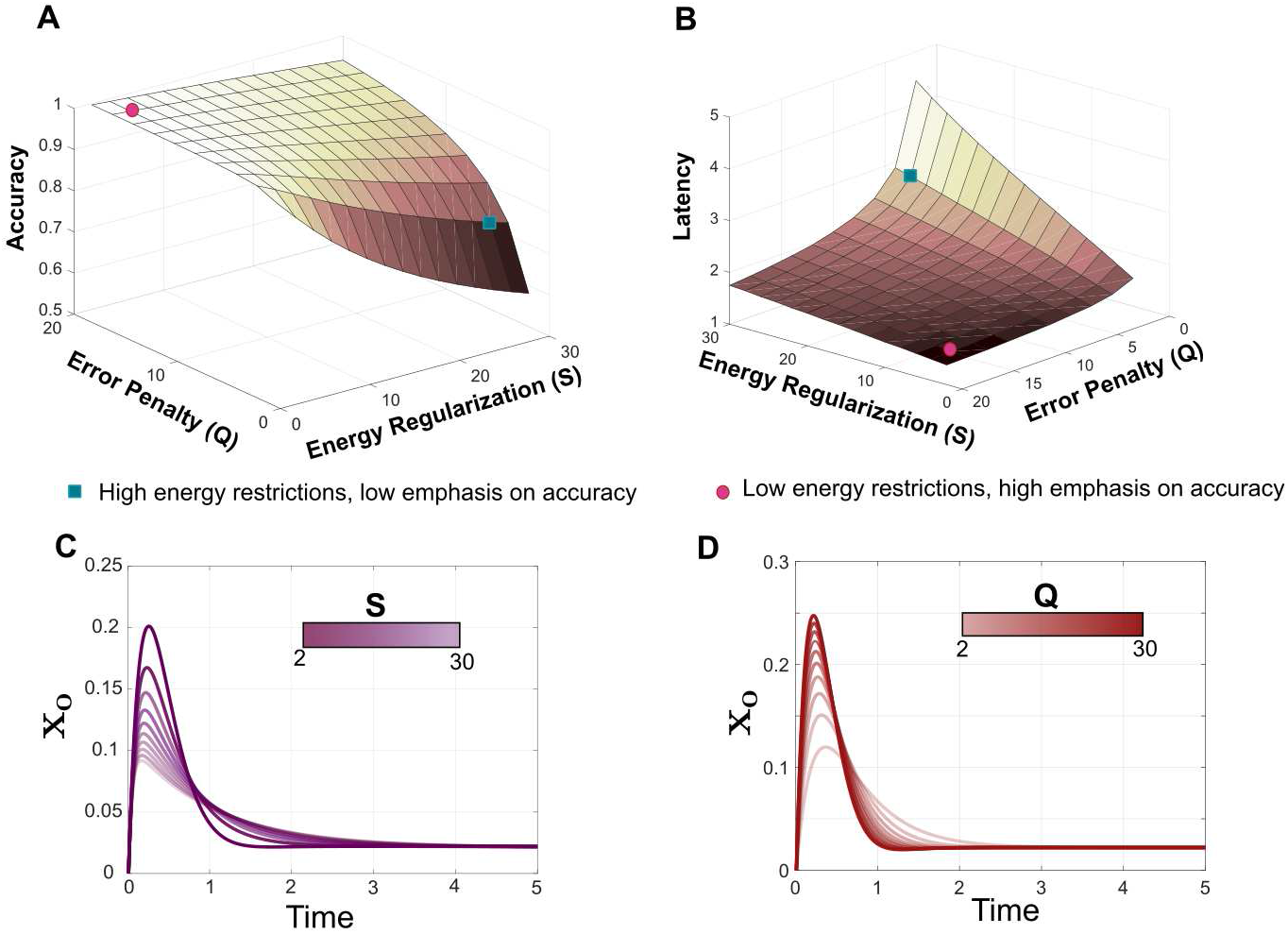
**A.** Accuracy is plotted as a function of error penalty matrix ***Q*** and energy regularization matrix ***S***. **B.** Latency of forming a reasonably accurate representation is plotted as a function of error penalty matrix ***Q*** and energy regularization matrix ***S***. (Note that the axis directions in **A** and **B** are reversed for better visualization of surface plots.) **C.** Response motifs across choice of energy regularization matrix ***S***. With a higher penalty on energy, the amplitude of the overshoot decreases. **D.** Response motifs across choice of error penalty matrix ***Q***. With highly scaled error penalty, the systems produce a sharp overshoot followed by a prompt return to baseline.

## Discussion

In this work we have proposed a mechanistic model for sensory detection that identifies the functional significance of spatiotemporal motifs observed ubiquitously across species in the context of olfaction. Previous studies on this topic (Stopfer et al., 2003; Bathellier et al., 2008; Raman et al., 2010; Mazor and Laurent, 2005; Saha et al., 2013b, 2017) have used post-hoc analysis to surmise the role of the aforementioned responses. In contrast, we present here a normative modeling perspective: we formally optimize PN response dynamics to achieve input separation in an intermediate latent space in an energy efficient manner.

The key contributions of this paper are summarized thus:

1. We showed that sensory onset and offset responses can be explained by a latent decoding objective function and associated mathematical optimization thereof.
2. We showed that a biologically plausible network architecture is capable of producing the optimal responses elucidated by 1.
3. We showed that this optimization leads to several emergent predictions regarding the mapping from stimulus to latent representation for more complex stimuli.

### Normative interpretation of PN response dynamics

We adopted a top-down approach and formulated a representation tracking problem wherein PNs must fluidly and efficiently encode the stimulus such that it generates a robust and coherent latent representation for downstream processes. It turned out that in order to achieve this objective, the optimal PN dynamics exhibited two distinct overshoots above baseline activity, one on onset of stimulus and the other on withdrawal (see Fig. 3). The analytical responses also exhibited distinct geometric orientations during stimulus on and off phases, analogous to what has been reported *in vivo* (see Figure 5). It is worth noting here that the optimization problem we have solved amounts to the linear quadratic regulator problem, a classical construct in control theory (Anderson and Moore, 2007).

We have shown here that an excitatory-inhibitory network architecture with first order dynamics can implement the motifs generated by the optimization framework. We derived the connectivity weights between the second order PNs and an auxiliary population of LNs by means of an iterative optimization scheme (see Figure 2 and Appendix for details). Our results however indicate that there might be several plausible ways to mathematically specify such a network.

We probed our model further to highlight the relevance of certain response features that appear both in simulations and through experimentation. The tractable construct facilitates a comprehensive understanding through rigorous mathematical simulations for different ranges of parameters and model alternatives. We analyzed the quality of the latent space representations formed using metrics of accuracy and latency to interpret what benefits neural response features such as competitive inhibition, contrast enhancement etc. provide in detecting a stimulus. A key result of this study was that the constructed sensory networks were capable of distinguishing between highly similar inputs (see Figure 9). There is in fact no inherent tradeoff between accuracy(i.e., input separation) and speed, rather both are monotonic with respect to energetic considerations(see Figure 10). Several studies in literature have discussed these notions through experimental observations and data driven, bottom-up modeling approaches (Wehr and Laurent, 1999; Ito et al., 2008; Rabinovich et al., 2000; Saha et al., 2017; Chittka et al., 2009). Here we presented a top-down normative framework that began from the specification of a mathematical form for the decoding objective and systematically navigated the results to provide a functional interpretation of patterns observed *in vivo*.

### Normative model produces known sensory neural dynamics beyond onset and offset responses

From our high-level detection objective, our normative model predicted several complex neural computations observed in the early olfactory networks beyond onset and offset responses.

For instance, experimental studies (Mazor and Laurent, 2005; Galán et al., 2004) have established that the trajectory in the neural state space quickly converges to a stimulus specific ‘fixed-point’: the transient activity being crucial in driving the trajectories towards these fixed-point attractors. Persistence of weak response (tonic activity) after the initial strong onset response(phasic activity) emerged in the synthesized model due to interplay between the first and second terms in the objective function, namely, accuracy of latent representation and energy of sensory activity (see Figure 8). So, the model indeed provides an objective way to think about the need for these ‘fixed-point’ attractors.

Additionally, our model also provided a novel interpretation of contrast enhancement computation (Nizampatnam et al., 2018) known to exist at a population level in PNs. It turned out that by highlighting the uniqueness of the current stimulus while sensing stimuli in series with very small inter-stimulus intervals, the PNs ensure robustness of latent representation required for reliable functioning of downstream processes(see Figure 6).

To summarize, we found that our framework built only to ensure accurate representation tracking in a lowdimensional latent space, exhibits (emergent) observations reported in the activity of olfactory systems, thus providing a unified explanation for these activity patterns.

### Generality of the decoding model

The decoder we have used is a multivariate linear dynamical system driven by PN activity. Mathematically, this is similar to drift-diffusion type models that have been used to study high-level decision making (e.g., the two-alternative forced choice task, (Colman, 2015; Bogacz et al., 2006)). In addition to the obvious difference in the level of behavioral abstraction being considered, there are important differences in formulation between our work and these prior results. In particular, our framework caters to the need of representations being fluidly created then abolished: they must *persist* through the duration of the stimulus and *dissipate* quickly after its withdrawal. This departs from theoretical decision-making paradigms such as *‘Interrogation’* or *‘Free Response’* (Bogacz et al., 2006) where a subject must make a detection within a set amount of time, and where the decoding process essentially terminates at the time of decision. Indeed, there is no ‘threshold’ in our model, only a latent representation that is continuously evolving in its state space.

In the scenario where the number of excitation sources i.e., odors is very large(possibly infinite), we can continue to use this normative framework for analysis by choosing an appropriate **b** matrix. In such a case, we could formulate **b** so that each row contains weights corresponding to a member of an appropriate basis set. Then, it is possible to translate the infinite set of inputs to their corresponding latent representations by linear combination of the chosen basis functions. The exact weight distributions for such basis set is yet to be determined and remains a question for future study.

### Features not explained

There are a number of important caveats and limitations that should be pointed out regarding the interpretation of the normative model presented through a set of first order firing rate dynamics (Dayan and Abbott, 2001). In particular, our synthesized network has linear dynamics wherein neurons can assume both positive and negative firing rates. Such a formulation is acceptable if we envision the rate variables as a change relative to some positive baseline rate of activity. Harder to reconcile in the linear model is a lack of upper and lower bounds of firing rate. Solving the optimization problem with such constraints is a harder problem mathematically (Chachuat, 2007) that we have not yet resolved.

Our construction produces local neurons that are exclusively inhibitory in nature and they interact only via the excitatory neurons. However, there are reports of existence of excitatory local neurons in the antennal lobe of some insects (Shang et al., 2007; Masse et al., 2009; Assisi et al., 2012). The functional role of such excitatory LNs is not accounted for in our model. However, in the mathematical formulation, these excitatory components can easily be realized by relaxing the non-positivity constraint in the iterative optimization procedure (see Appendix for details). Our construction also produces very weak lateral inhibition directly between PNs (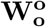, see Figure 2). Although there are no experimental evidence of existence of such PN-PN inhibitory synapses, fast GABAergic synapses (Bazhenov et al., 2001, 2005) are known to exist in the insect antennal lobe (between LN-LN and LN-PN) and thought to be functionally relevant for maintaining the synchrony of PN responses (Bazhenov et al., 2001). We theorize that the inhibitory synaptic weights between PNs produced by solving the reduced LQR problem (9) are in fact proxies for these fast inhibitory synapses that have not been explicitly considered in the synthesized network.

In the objective function (Eq. 2) we use a Euclidean distance to evaluate the quality of the latent representation as it tracks the incoming stimulation. However, downstream neurons might achieve input separation prior to higher level processing by more involved computations. Experimental studies identify spatial and temporal filtering, decorrelation etc. (Friedrich, 2013) as operations that could be plausibly executed by sensory neurons. In the present work we choose to use the well characterized *ℓ*_2_ norm as a surrogate for specialized computations that might occur *in vivo*. It is as yet an open question how incorporating such functions would impact our synthesis results.

Additionally, we have not considered any form of adaptation or habituation in our network, choosing to focus on a single exposure regime. The question of how the optimal weights come to be, e.g., via development or learning, is not considered herein.

Finally, our model makes no statements about the translation from latent representations to behavior. Behavior can exhibit nonlinear and sometimes paradoxical characteristics with respect to stimulus intensity, composition and context. Accounting for these factors within the optimization problem is highly intriguing, but a more complex formulation that is left for future study.

## Acknowledgements

This work has been supported by grants EF-1724218 (SC,BR), IIS-1453022 (BR), CMMI-1653589 (SC) from the National Science Foundation. ShiNung Ching holds a Career Award at the Scientific Interface from the Burroughs-Wellcome Fund.

## Appendix

## Reduction of optimization framework to Infinite Horizon Linear Quadradic Regulator problem

In this section show how our problem reduces to a known design framework in systems engineering. The optimization problem (OP) posed is:

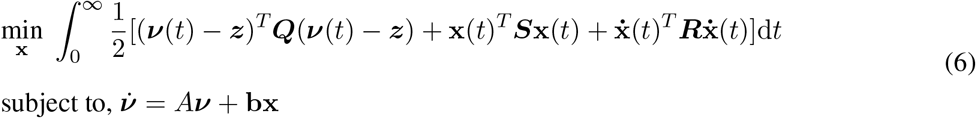

We denote 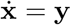. Thus, our OP can be rewritten as,

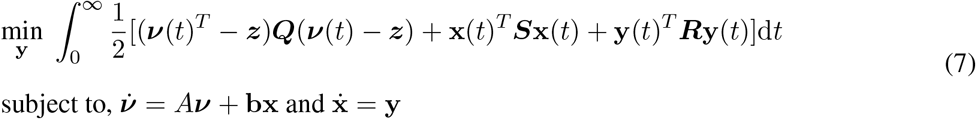

Now, we define **v**_*z*_ = [***ν***^*T*^, **x**^*T*^, **z**^*T*^]^*T*^ . Also, we assume that the abstract variable ***z*** remains constant as long as the same stimulus conditions prevail i.e., 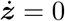. Therefore,

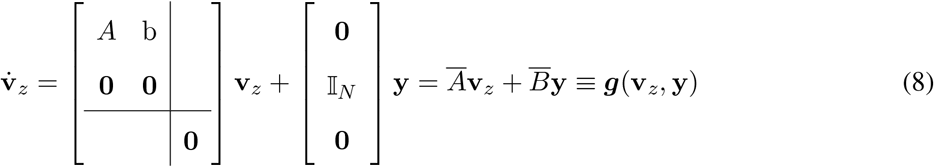

With this definition, (7) reduces to:

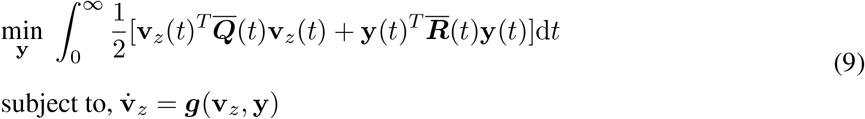

where, 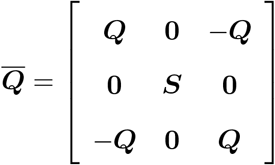 and 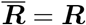.

The transformed OP (9) amounts to an Infinite Horizon Linear Quadratic Control problem (Anderson and Moore, 2007; Boyd and Barratt, 1991), a classical state-space design framework for finding exogenous controls for a dynamical system. In our context, the ‘controls’ are the activity of the PNs and the task is to realize these ‘controls’ by means of a dynamical network.

## Solving the optimization problem

As the matrices ***Q***, ***S*** and ***R*** in (7) are positive definite, it is straightforward to establish that 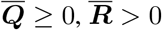 and, hence, the problem (9) has a unique solution (Anderson and Moore, 2007). Indeed, the solution to (9) with initial conditions 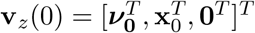. It is given by,

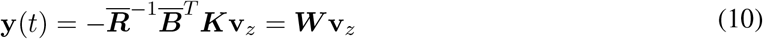

where ***K*** is the solution of the Algebraic Riccati Equation (ARE), given as follows:

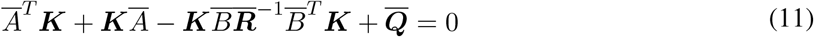

Therefore, we can now write,

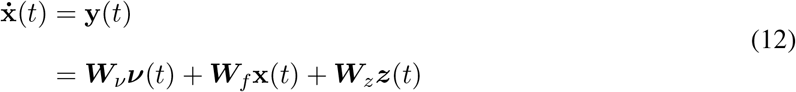

where, ***W*** = [***W***_*ν*_ : ***W***_*f*_ : ***W***_*z*_] and ***W***_*ν*_ ∈ ℝ^*n×m*^, ***W***_*f*_ ∈ ℝ^*n×n*^ and ***W***_*z*_ ∈ ℝ^*n×m*^.

The excitation to the PN layer arrives from first-level ORNs. We denote this excitation as **r**(*t*). In this model therefore,

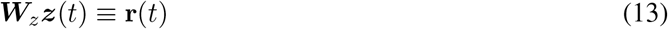

It is important to note that the model does not predict ORN dynamics or connectivity per se, only that ***r***(*t*) is provided to the network the PNs. There may be multiple ways to synthesize an ORN network that realizes this transformation, and this question is left for future work.

Again, recall,

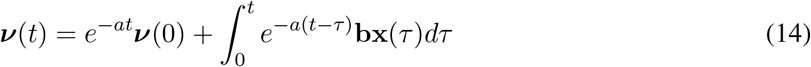

We initialize the decoder to a zero initial state i.e., ***ν***(0) = **0**. Thus, (14) can be simplified to,

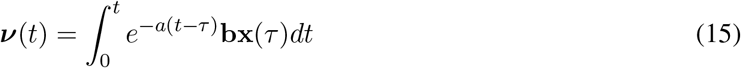

Using (14) and (13) in (12) we obtain,

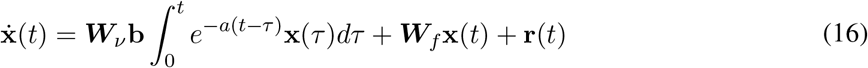

Eq. (16) represents the optimal PN dynamics that converts a peripheral stimulus (arriving via **r**(*t*)) to an intermediate representation for further processing by higher brain regions.

A similar problem can be posited when the latent decoder is a noisy one or when the neurons are associated with inherent background noise. In such a model the objective function is written as follows:

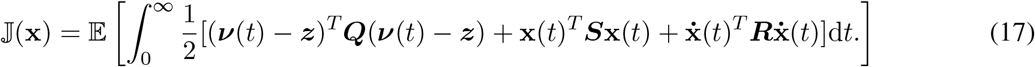

The reduction technique for this stochastic problem is similar to what has been shown above (Boyd and Barratt, 1991) and the solution of the Algebraic Riccati Equation(ARE) for the Stochastic LQR is identical to what we have derived here except for the addition of a noise term ***η***(*t*)(see Eq. (18) below).

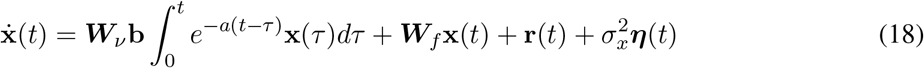

We have used this stochastic version to generate figures depicting PC trajectories and correlation of ensemble response as in Fig. 5C and Fig. 8B.

## Extracting a first order network to realize the optimal solution

The primary step in extracting a first order network from (16) is to break out the slow integration into a separate population of *n_i_* auxiliary neurons, i.e.,

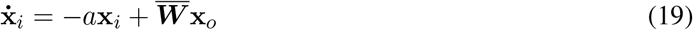

where 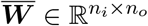 and *n_o_* is the number of PNs.

Thus, the function of the matrix 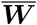 is to compute a weighted sum of the PN activity and propagate it to the auxiliary neurons. Using (19) in (16), we can write:

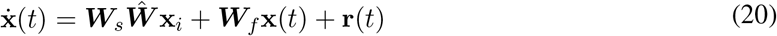

where ***W***_s_ ≡ ***W***_*ν*_**b** and 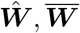 must satisfy the condition 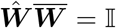. It has been observed *in vivo* that the number of neurons in the auxiliary population is less than the number of PNs, i.e., *n_i_ < n_o_* (Laurent, 1996). Due to this biophysical constraint 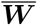 is a wide matrix and the matrix decomposition problem 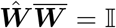 is ill-posed and might have no solution or an infinite number of solutions. Instead of solving this underdeter-mined set of equations for an arbitrary choice of 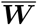, we reformulate the problem in an iterative regularized optimization framework (Shlizerman et al., 2014) (see Algorithm 1 in Parameter Selection below).

The synthesized first order network is therefore,

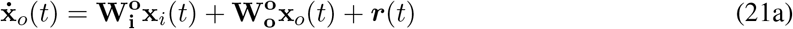

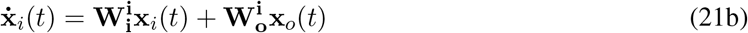

The matrices connecting different neural populations in this network are:

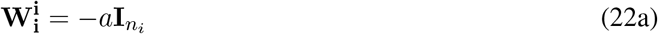

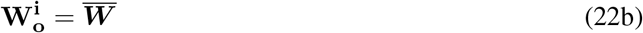

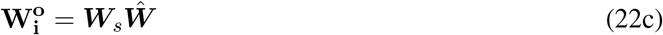

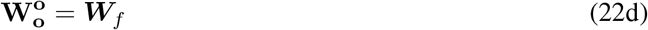

## Parameter selection

With the exception of Figure 7, we chose the decoder parameter as *a* = 0.25. The penalty matrices ***Q***, ***S***, ***R*** are all scaled identity matrices. The result of Figure 4 in particular resulted from ***Q*** = 10**I**_*m*_, ***S*** = 2**I**_*n*_, ***R*** = 0.2**I**_*n*_. Also, unless mentioned otherwise we have chosen *n* = 41 (number of neural units driving the decoder) and *m* = 2 (dimension of the latent space). For simulations using the stochastic model we have considered ***η***(*t*) to be unit variance Gaussian noise with *σ_x_* = 0.05.

The choice of matrices 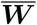 and 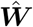 are determined by an iterative optimization scheme executed as: ‖.‖_2_ here is the Frobenius matrix norm and 0 < λ_1_ < 1, 0 < λ_2_ < 1 are coefficients of regularization. We initialize the algorithm with an arbitrary wiring for 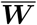 (eg: positive random block matrix) such that its pseudo-inverse 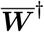 exists. The initial choice for 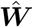 is 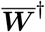, as the pseudoinverse is the best fit for a least squares solution. However, choosing the pseudoinverse for 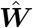 does not guarantee that 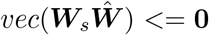. Thus, we need the iterative framework wherein we find a pair of matrices 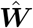 and 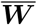 such that the resultant network realizes the optimal motifs and the biophysical constraints are satisfied. The optimization problem is solved using the disciplined convex optimization package CVX implemented in MATLAB (Grant et al., 2008). If the iterative optimization framework 1 returns a solution, then we can infer that the derived network architecture that is capable of producing the observed response motifs in presence of exogenous excitation. It is worthwhile to point out here that the motifs for LNs indeed depend upon the obtained 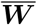 matrix as it encapsulates how each PN forms connections with the putative LNs. However, the iterative optimization scheme ensures that the phasic/tonic motif of PNs is preserved by generating inhibitory connections from LNs to PNs (since 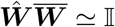).

**Algorithm 1:**
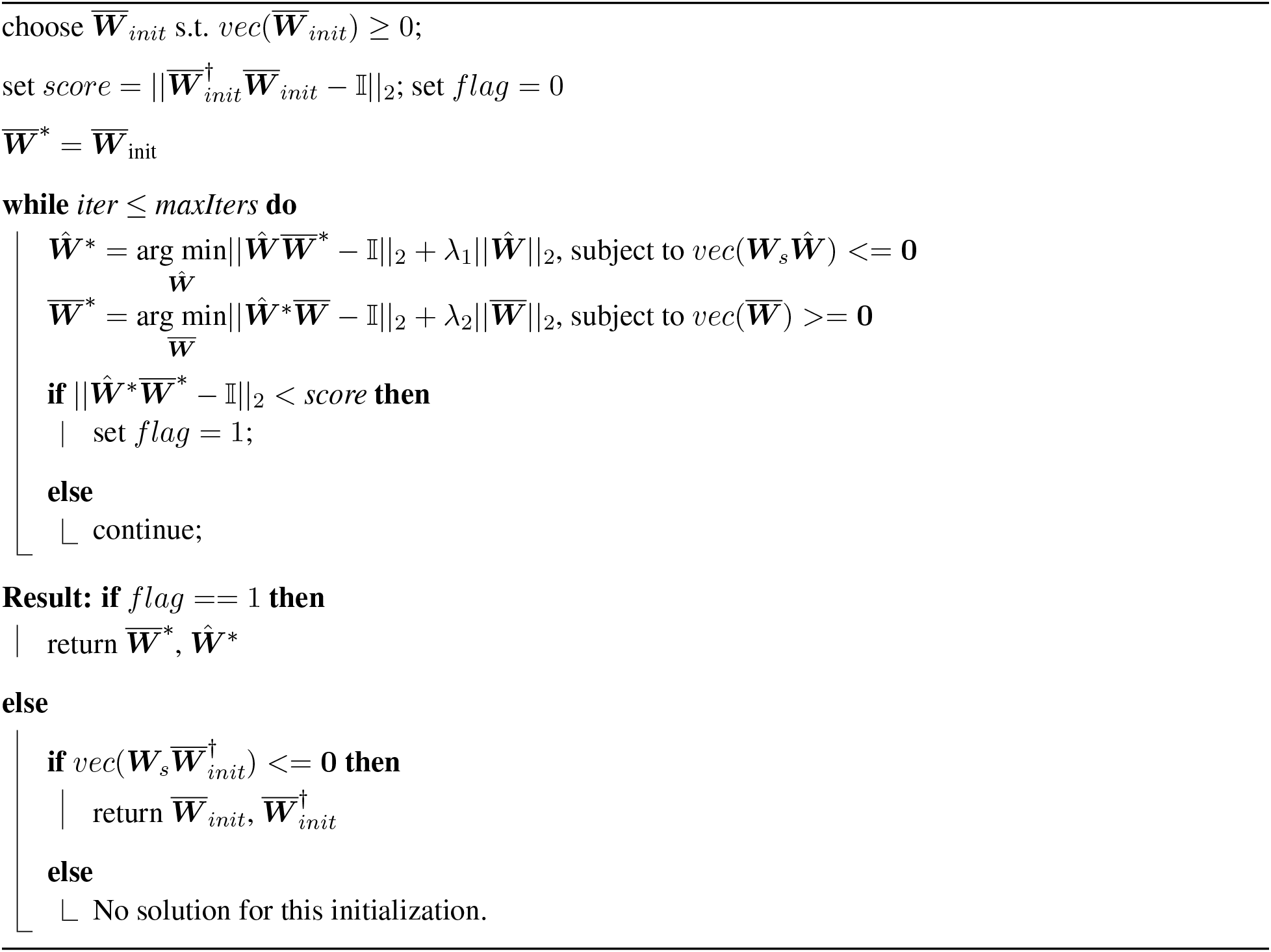

## Higher dimensional generalization of the latent space

In this section we address how can the latent space be generalized to higher dimensions so that a large number of odor inputs can be represented. Here, we consider six odors (blue/magenta/red/olive/green/cyan) that map to a three dimensional high-level feature space (see Figure 11A, B) through matrix **b**. Similar to the two dimensional case, neural activity causes accrued evidence ***ν***(*t*) towards track ***z*** in the latent space (see Figure 11C). Through this simulation, we find out that through choice of an appropriate tuning matrix it is possible to represent a relatively large number of odors in a low dimensional latent space.

**Figure 11:**
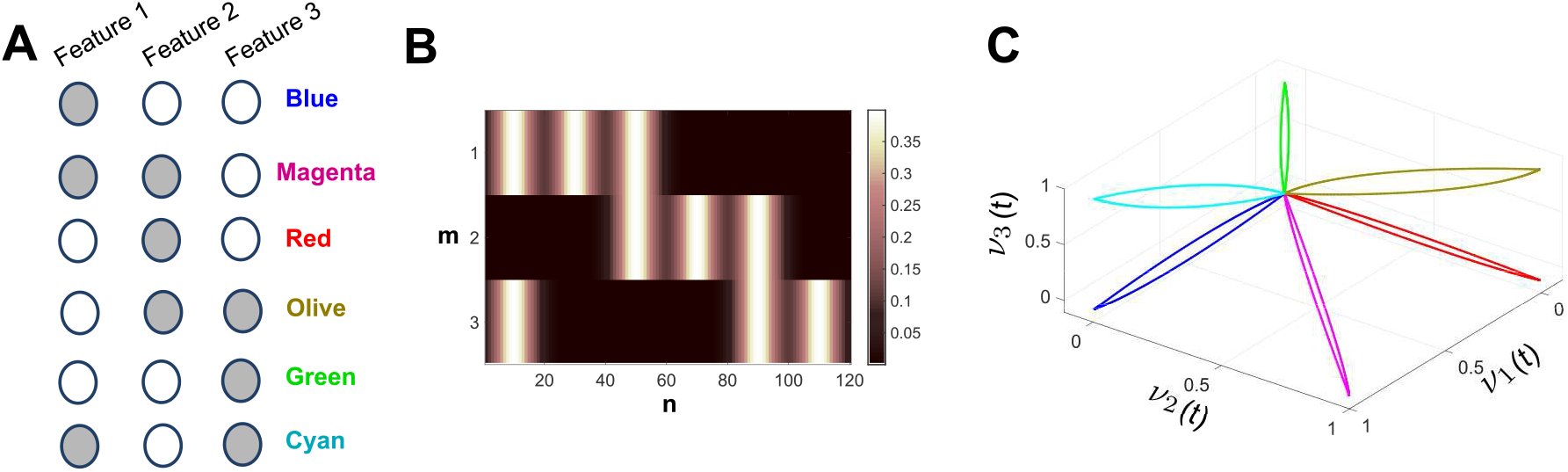
**A.** Odors and their respective mapping to the high level feature-space. **B. b** ∈ ℝ^*m×n*^ weighs the contribution of each neuron to each feature. **C.** Latent space trajectory for each of the six representative odors.

## Accuracy, Similarity and Latency

In Figure 9 and 10 we performed two quantification of the decoding performance on the presumption of a discrete pulsatile stimulus.

- **Accuracy**: Accuracy quantifies the deviation of the latent space representation produced by PN activity from the true representation. In Figure 9 this quantification was made at the end of the stimulus period, given mathematically by:

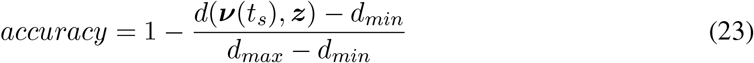

where, *d*(*u, v*) is a distant metric between the points u and v and *t_s_* is the time instant when the pulse in withdrawn. We use a Euclidean metric for distance in this case.
- **Similarity**: Another way of evaluating the quality of the latent representation produced by neural activity is by measuring the cosine of the angle between ***ν***(*t*) at time *t* = *t_s_* and ***z***. This can be interpreted as computing the similarity between the latent representation at the end of stimulus period and the desired nominal representation and is given mathematically as:

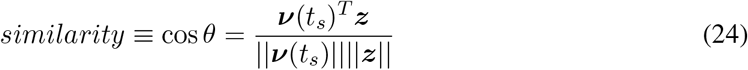
- **Latency**: Latency quantifies the time required by the synthesized network model to provide an accurate representation of the stimulus. It is computed as the time necessary to reach 1 − ϵ of the nominal representation ***z***. In Figure 10 we used ϵ = 0.2.

**Figure 12:**
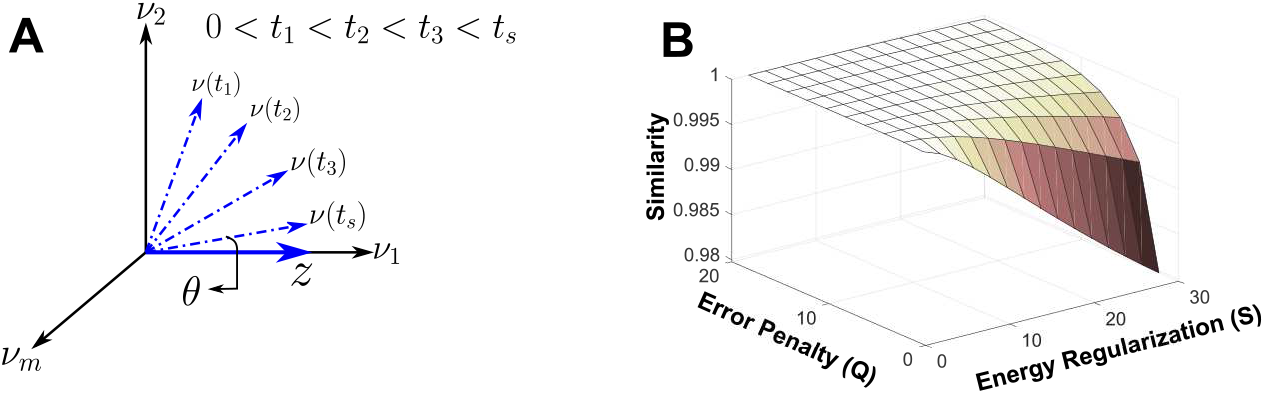
**A.** Schematic representation of evolution of latent representation and measurements relevant to computing quality of latent representation in terms of a cosine metric. **B.** Similarity is plotted as a function of error penalty matrix ***Q*** and energy regularization matrix ***S*** with the axis for error penalty ***Q*** reversed for better visualization.

## Dimensionality reduction analysis for simulated PN response

Similar to methods in (Saha et al., 2017), we applied Principal Component Analysis (PCA) on our simulated PN response data in order to visualize the *n*-dimensional time varying activity of PNs. Each exposure to a stimulus of given identity continues for *τ* seconds, followed by a period of no excitation of equal duration.As we synthesized PN response at 10ms intervals, we generated a time-series data matrix of dimensions *n ×* Γ (where 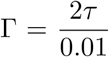) for each stimulus encounter. We concatenated such data matrices obtained for different odorants in order to compare and contrast the trajectories evoked. The resulting matrix was used to generate a *n × n* response covariance matrix.

In order to visualize the neural activity during on and off response phases, a time window comprising of activity at stimulus onset and withdrawal is selected (4s on followed by 4s off). This high-dimensional time-varying data is then projected along the first three eigen-vectors of the response covariance matrix. Low dimensional data points in consecutive time instances were connected to construct trajectories in the three dimensional space. On the other hand, to compare the activities invoked by red/blue stimulus in on and off phases, a 4s window comprising of either on or off response is used.

